# Uncovering temporally sensitive targeting motifs for traumatic brain injury via phage display

**DOI:** 10.1101/2020.06.16.155325

**Authors:** Briana I. Martinez, Gergey Alzaem Mousa, Kiera Fleck, Tara MacCulloch, Chris W. Diehnelt, Nicholas Stephanopoulos, Sarah E. Stabenfeldt

## Abstract

The heterogeneous injury pathophysiology of traumatic brain injury (TBI) is a barrier to developing highly sensitive and specific diagnostic tools. Embracing neural injury complexity is critical for the development and advancement of diagnostics and therapeutics. The current study employs a unique discovery pipeline to identify targeting motifs that recognize specific phases of TBI pathology. This pipeline entails *in vivo* biopanning with a domain antibody (dAb) phage display library, next generation sequencing (NGS) analysis, and peptide synthesis. Here, we identify targeting motifs based on the HCDR3 structure of dAbs for acute (1 day) and subacute (7 days) post-injury timepoints using a mouse controlled cortical impact model. Their bioreactivity was validated using immunohistochemistry and candidate target epitopes were identified via immunoprecipitation-mass spectrometry. The acute targeting motif recognizes targets associated with metabolic and mitochondrial dysfunction whereas the subacute motif was largely associated with neurodegenerative processes. This phage display biomarker discovery pipeline for TBI successfully achieved discovery of temporally specific TBI targeting motif/epitope pairs that will advance the TBI diagnostics and therapeutics.

## INTRODUCTION

Traumatic brain injury (TBI) affects an estimated 1.7 million people in the United States each year and is a leading cause of death and disability for children and young adults in industrialized countries [1–3]. Individuals who experience TBI are more likely to develop cognitive and behavioral deficits, as well as physical conditions such as inhibited motor coordination and balance [4]. These individuals are also more susceptible to acquiring neurodegenerative diseases than the non-injured population [5, 6]. Treatment costs of TBI are estimated at $76.5 billion annually in the United States alone [7], making TBI a great economic burden and public concern.

TBI is characterized not by a singular event, but a cascade of two separate injury phases. The initial insult disrupts the blood-brain barrier (BBB) and causes necrosis, tissue deformation, and cell shearing [8]. This primary injury then catalyzes the secondary injury cascade, leading to an increase of inflammatory cytokines, mitochondrial damage, ischemia, and cell death [8, 9]. This pathology may persist for hours to months after the initial insult, introducing temporal complexity to the injured neural milieu [10]. Unfortunately, the molecular and cellular mechanisms of injury progression are complex and have yet to be fully elucidated. Consequently, this complexity impacts the development of accurate diagnosis and treatment options.

Biomarkers, objective signatures of injury, can inform and facilitate development of sensitive and specific theranostic devices. For example, a quantifiable biomarker can provide insight on the severity of a patient’s injury or be utilized to assess treatment efficiency. Biomarkers can also inform the design of targeted therapeutic agents. However, the temporal and dynamic evolution of TBI is a barrier to developing sensitive theranostic devices. A biomarker identified acutely after injury may not carry the same diagnostic or therapeutic value at more chronic timepoints and vice versa. Accordingly, the identification and characterization of temporal TBI biomarkers is critical for improving patient care. Molecular-based biomarker discoveries are often facilitated with “top-down” approaches, where known involvement in the condition is paramount to classification as a candidate biomarker. Advances in neuroproteomics have been applied to TBI biomarker discovery to support “bottom-up” approaches [11, 12]. This method takes advantage of the heterogeneous injury environment by fractionating lysate and analyzing global protein expression via mass spectrometry to identify proteins with differential expression after injury. Several studies have successfully analyzed the proteome of rodent brain tissue after experimental TBI to uncover hundreds of differentially expressed proteins that have the potential to be candidate biomarkers of neural injury [11, 13, 14]. However, neuroproteomics approaches often yield a large volume of data, leading to time intensive analysis and difficulties selecting candidates for clinical use [11, 12].

Phage display, a powerful screening technique to uncover protein-protein interactions, has been applied to biomarker discovery in various cancers and more recently, neurological conditions such as Alzheimer’s disease and stroke [15–19]. This technique utilizes libraries of biological motif-displaying bacteriophages that are then screened against target antigens. Bound bacteriophage are then collected and amplified to use in a subsequent biopanning round. This process is repeated to enrich the population of motifs that have strong affinity to the target antigen. Several display systems can be applied to biomarker discovery, but the domain antibody fragment (dAb)-based display systems in particular hold distinct advantages. These fragments represent the variable heavy (V_H_) domain of a full-length antibody and contain three heavy complementarity determining regions (HCDRs; 1, 2, and 3) to facilitate stability. The HCDR3 specifically is the primary determinant of antigen binding [20–22]. The small size of the dAb structure (12-15 kDa) and ability to isolate proteins specific to brain vasculature make them ideal for interacting with the neural milieu *in vivo* [23, 24]. Despite these characteristics, implementation of dAb phage display to elucidate temporal mechanisms of TBI has yet to reach its full potential.

The difficulty of utilizing antibody fragment systems lies in production of individual candidate fragments for bioreactivity validation, specifically when taking advantage of next generation sequencing (NGS) analysis of phage libraries [25]. With classical approaches, phagemid DNA is directly isolated for validation while deep sequencing approaches rely on fragment assembly and recombinant protein production techniques [25, 26]. Methodology to produce dAbs is time consuming and resource intensive due to difficulties maintaining stability after purification and degradation of affinity tags [27, 28]. To address these limitations “synthetic antibody”-type constructs have been developed by taking advantage of the antigen binding qualities of the dAb HCDR3 [29–32]. These constructs are functional at an even smaller scale, as they are solely derived from the HCDR3 sequence and modified for stability. HCDR3 constructs have been shown to mimic the binding specificity and capacity of full-length antibodies for factors such as platelet aggregation and HIV-1 promoter at a fraction of the size [30, 32] and demonstrate the ability to be integrated with nanoparticle engineering to facilitate protein-like functionality in the nanoparticle [29].

Here, we use *in vivo* phage display biopanning to take advantage of the heterogeneous injury milieu and dAbs that specifically target injured neural tissue at 1 and 7 days post-injury (dpi) in a mouse model of controlled cortical impact (CCI). HCDR3 constructs were developed based on next generation sequence (NGS) and bioinformatic analysis of dAb phage populations and spatiotemporal specificity was validated via immunohistochemical analysis. Through immunoprecipitation-mass spectrometry (IP-MS) with the HCDR3 constructs, we identified several potential targets associated with metabolic dysfunction and neurodegenerative processes expressed at the acute and subacute timepoints, respectively.

## MATERIALS AND METHODS

### Controlled cortical impact

All experiments were approved by the Arizona State University Institutional Animal Care and Use Committee (IACUC).

Eight-weeks-old male and female C57Bl/6 mice (Charles River) were assigned to four experimental groups; acute (sacrificed 1 dpi), subacute (7 dpi), chronic (21 dpi) and sham (craniotomy with no injury, sacrificed 1 day post-procedure). Mice were further divided for each experimental assay: biopanning, immunohistochemistry (IHC), or IP-MS whereby each experimental condition and analysis technique had a range of n = 6-10. Briefly, mice were anesthetized with isoflurane (3% induction, 1.5% maintenance and secured on a stereotaxic frame (Leica Microsystems, Wetzlar, Germany). A 3 mm craniotomy (−1 AP mm bregma, 1.5 lateral to midline) was performed to accommodate a 2 mm diameter, 1 mm deep impact to the frontoparietal cortex at 6 m/s velocity and 100 ms duration. Surgical area was sutured, then analgesics (0.05mg/kg buprenorphine) and saline were subcutaneously administered. Mice were placed in single housed cages and monitored during recovery.

### *In Vivo* Biopanning

A human dAb library (Source Bioscience, Nottingham, UK) was prepared with hyperphage (Progen, Heidelberg, Germany) as described in the manufacturers protocols [33, 34]. At 1,7, or 21 dpi, the parent phage library (10^12^-10^14^ CFU in 100 µL saline) was administered via retro-orbital injection. Phage circulated for 10 minutes before animals were euthanized via pentobarbital solution overdose (150 mg/kg intraperitoneal injection). Non-specific phage were cleared after thoracotomy and transcardial perfusion with 0.1M phosphate buffer, pH 7.4. The ipsilateral (injured) and contralateral hemisphere of the brain were extracted and dissected, in addition to the peripheral organs – heart and spleen. Simultaneously, this procedure was repeated in sham animals. Immediately, tissues were weighed, homogenized, pooled (n=3/biopanning round), and mechanically homogenized in chilled phosphate buffer with protease inhibitors. Trypsin was added to the homogenate to elute binding phage from tissue. Phage concentration determined by colony forming units (CFU) of tissue elutions were quantified by bacteria titers (TG1 E. coli). Titers were completed after each round to confirm distribution across tissues. Eluted phage were amplified with TG1 E. coli and stored under −80°C conditions. Between biopanning rounds, phage DNA were isolated (QIAprep Spin Miniprep; Qiagen, Valencia, CA) and analyzed for sequence convergence by the DNASU Sequencing Core (Tempe, AZ).

For the second biopanning cycle, the eluted phage from the ipsilateral brain hemisphere were amplified and purified to serve as the phage population for the second biopanning round. Injection, tissue preparation, phage elution, amplification, and storage were completed as stated previously. A stock library from the manufacturer was amplified without a screening target to serve as a propagation library control to prevent selection of non-specific, parasitic sequences.

### Next Generation Sequencing and Analysis

Preparation of phage dAb libraries for deep sequencing was completed following the Illumina Nextera XT amplicon sequencing protocol (Illumina, San Diego, CA). Briefly, amplicons were created with a single PCR step and Illumina-compatible indexes were added to each sample with a second PCR cycle. Phage libraries were sequenced with primers including Illumina-specific barcodes (Supplementary Table 1) via Illumina MiSeq 2 x 300 bp module by the DNASU Next Generation Sequencing Core at ASU Biodesign Institute (Tempe, AZ).

Paired end sequences were stitched with Fast Length Adjustment of Short reads (FLASH) [35], using a minimum and maximum overlap of 10 and 200 bp respectively. The (HCDR3) sequence of each dAb was extracted in Bioconductor for R [36] by subsetting between dAb frameworks 3 and 4. Mutated HCDR3 sequences were excluded from analysis by filtering for sequence length. Raw reads and normalized reads per million (RPMs) were retrieved with the FASTAptamer Toolkit [37]. HCDR3 sequences in injury groups that were enriched through biopanning (i.e. increase of reads from round 1 to round 2) were selected from each library. Enriched sequences were then compared with peripheral tissue (spleen and heart) and propagation control libraries to ensure final selection of HCDR3s that were specific to injured neural tissue libraries. Further, selected sequences were compared against other injury timepoints (i.e., sequences selected from acute injury were compared with sequences from the subacute injury group) using z-scores to enhance temporal specificity for each sequence selection. Top HCDR3s were selected for antibody-mimetic production and further validation based on the following criteria: 1. frequency, 2. fold enrichment values, and 3. specificity to neural injury at the distinct biopanning timepoints.

### Biotinylated HCDR3 constructs

HCDR3 peptides for injury and control groups were synthesized with N- and C-terminal Cys for preparation of constrained peptide constructs (WatsonBio, Houston, TX). Peptides were then conjugated to a bivalent peptide linker [38] that utilized thiol-bromoacetamide conjugation to constrain the peptides via the N- and C-terminal Cys. The peptide linker contained a C-terminal biotin. Each reaction was analyzed by matrix assisted laser desorption/ionization to confirm presence of peptide-scaffold conjugate (Supplementary Figure 1). Constructs were purified via high performance liquid chromatography (HPLC) collection at 214 nm for downstream analyses.

### Immunohistochemistry

Mice were subject to CCI or sham (n=3 per group/sex) as described previously and perfused with 0.1 M phosphate buffer and 4% paraformaldehyde at designated timepoints. Brains were fixed overnight in 4% paraformaldehyde at 4°C followed by immersion in 15% sucrose and then 30% sucrose. Brains were flash frozen on dry ice in optimal cutting temperature medium and stored at −80°C. Samples were sectioned coronally at 20 µm thickness. After incubation with excess streptavidin and biotin to block endogenous biotin (Endogenous Biotin Blocking Kit; Thermo Fisher Scientific, Waltham, MA), 5 µM of biotinylated HCDR3-construct was incubated on tissue overnight at 4°C. Simultaneously, control sections were incubated with control HCDR3-construct or 1X PBS. Tissue sections were washed in 1X PBS, then HCDR3-construct samples were incubated with Alexa Fluor 555 streptavidin at room temperature for 2 hours, followed by 1X PBS washes. Tissue sections were subject to DAPI incubation for 5 minutes. Sections were visualized using fluorescence microscopy, and 20x magnification tile scans were prepared for processing using ImageJ software. Threshold values of control slices were used to quantify fraction of HCDR3 construct positive staining. Analyzed area was approximately 1500 µm x 1500 µm.

### Immunoprecipitation-mass spectrometry

CCI and sham surgeries were completed as described previously. Mice were sacrificed at 1 or 7 dpi (n=3/group) via transcardial perfusion with phosphate buffer, pH 7.4. The ipsilateral hemisphere of the brain was immediately dissected and homogenized in chilled lysis buffer (1X PBS, 1% Triton, protease inhibitor cocktail). Protein concentration was quantified with the Pierce BCA Protein Assay Kit (Thermo Fisher).

Streptavidin-coupled Dynabeads (Thermo Fisher) were washed with 0.1% Tween in 1X PBS and incubated with 1 mg/mL tissue lysate for 1 hour at room temperature. Pre-cleared lysate was collected after separation from magnetic beads and incubated with designated HCDR3-constructs rotating overnight at 4°C to form the immune complex.The immune complex was incubated with streptavidin-coupled Dynabeads for 1 hour at room temperature and beads were then washed 3 times with chilled lysis buffer. For A2, captured antigens were eluted by heating sample at 95°C with SDS PAGE running buffer. Samples were run on a stain-free 12% SDS-PAGE gel. Protein bands were excised and processed by the ASU Biodesign Mass Spectrometry Facility for liquid chromatography-mass spectrometry (LC-MS) analysis using the Thermo Orbitrap Fusion Lumos (Thermo Fisher). For SA1, the same immunoprecipitation and wash procedure were completed, and antigens were eluted directly from beads with 0.2% Rapigest for LC-MS analysis.

UniProt IDs of identified proteins were uploaded to the PANTHER (Protein Analysis Through Evolutionary Relationships) classification system [39] and searched against the Mus musculus reference database. Ontological assessments to characterize cellular localization, molecular function, biological processes, and pathways were conducted with PANTHER Overrepresentation test with the GO Ontology database (released 2019-10-08).

### Statistics

For NGS analysis, raw counts were first normalized to reads per million (RPM) to account for library differences. A normalized z-score was then used as a threshold to identify dAbs that were highly represented and specific to their distinct injury timepoint. Selected dAbs were then screened for enrichment factor and individual frequency. For IHC, fluorescence percentage per area was conducted with ordinary one-way ANOVA followed by Tukey’s test for multiple comparisons; statistical significance was determined as p < 0.05. Identified proteins that met the false discovery rate (FDR) threshold of < 0.01 were used in all ontological assessments to categorize biological processes and candidate pathways.

## RESULTS

### dAb phage bind to injured brain tissue *in vivo*

A dAb phage library was intravenously injected into CCI injured mice at 1, 7, and 21 dpi (Figure 1). Phage accumulation was analyzed through titer analysis to confirm that the phage library was given sufficient time to bind to target tissues. Titers determined that phage accumulated in all extracted tissues, with the spleen having the highest total CFU/g tissue of 1.05 x 10^7^ (Supplementary Table 2). Up to 1.21 x 10^6^ CFU/g tissue were recovered from neural tissue of each cohort through trypsinization, including sham controls. An increase in ipsilateral hemisphere-binding phage was observed comparing biopanning round 1 to round 2 for both the acute and subacute timepoints (increases of 28 and 37% respectively), indicating successful enrichment of affinity binders to target tissue (Figure 2b,c). The percent of CFU bound to chronic injured and sham tissue were similar between biopanning rounds (Figure 2a,d).

**Figure 1:**
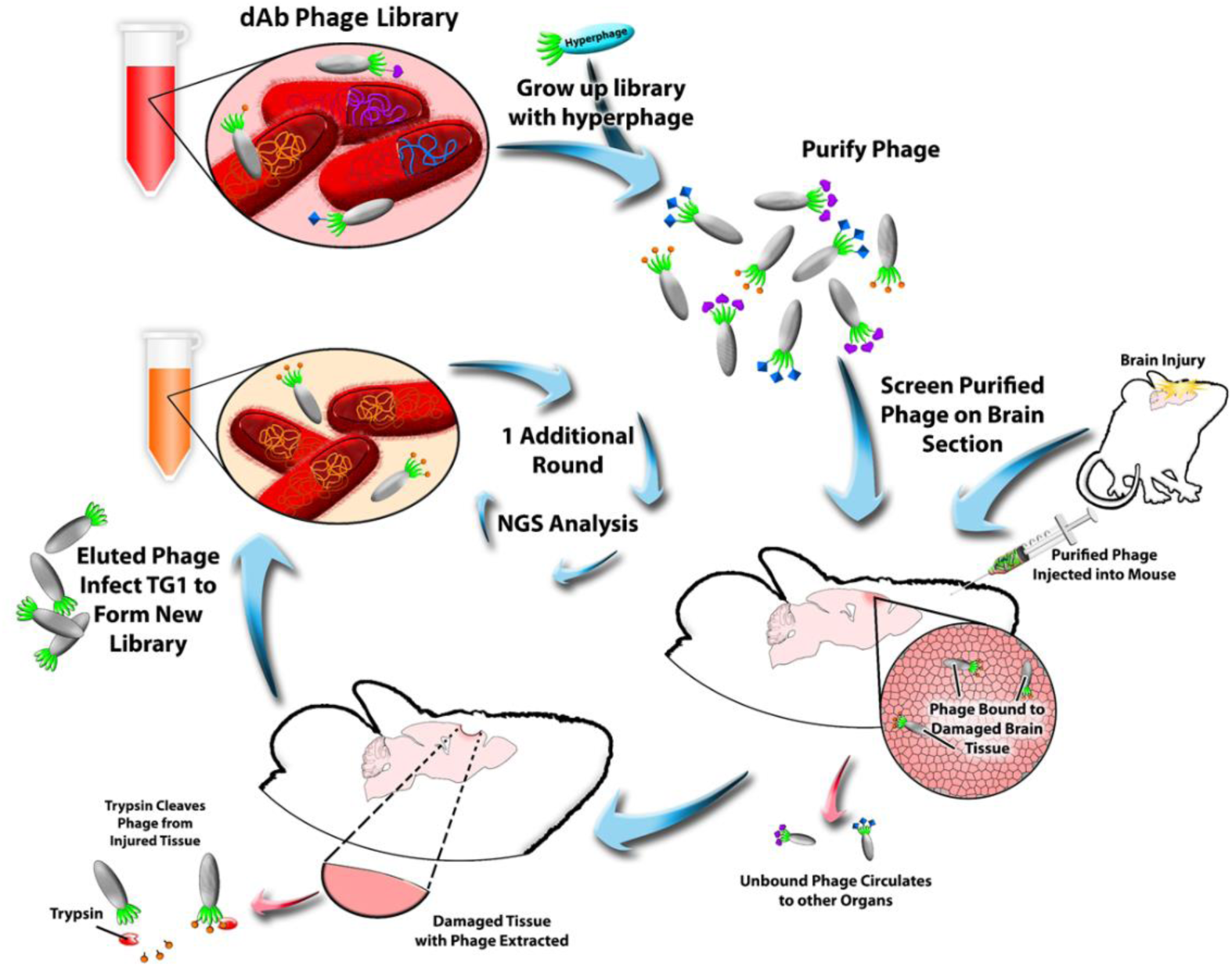
Schematic of phage display biopanning. A dAb phage parent library is produced and purified, then intravenously injected into a mouse that has had a controlled cortical impact (CCI) at a distinct timepoint (1, 7, or 21 dpi) or a sham injury (sacrificed 1 day post procedure). Tissue are extracted, lysed, and trypsinized to cleave phage from tissue. The phage library from the ipsilateral hemisphere is then amplified with TG1 E. coli and applied in the final round of biopanning. Recovered phage are then analyzed using NGS.

**Figure 2:**
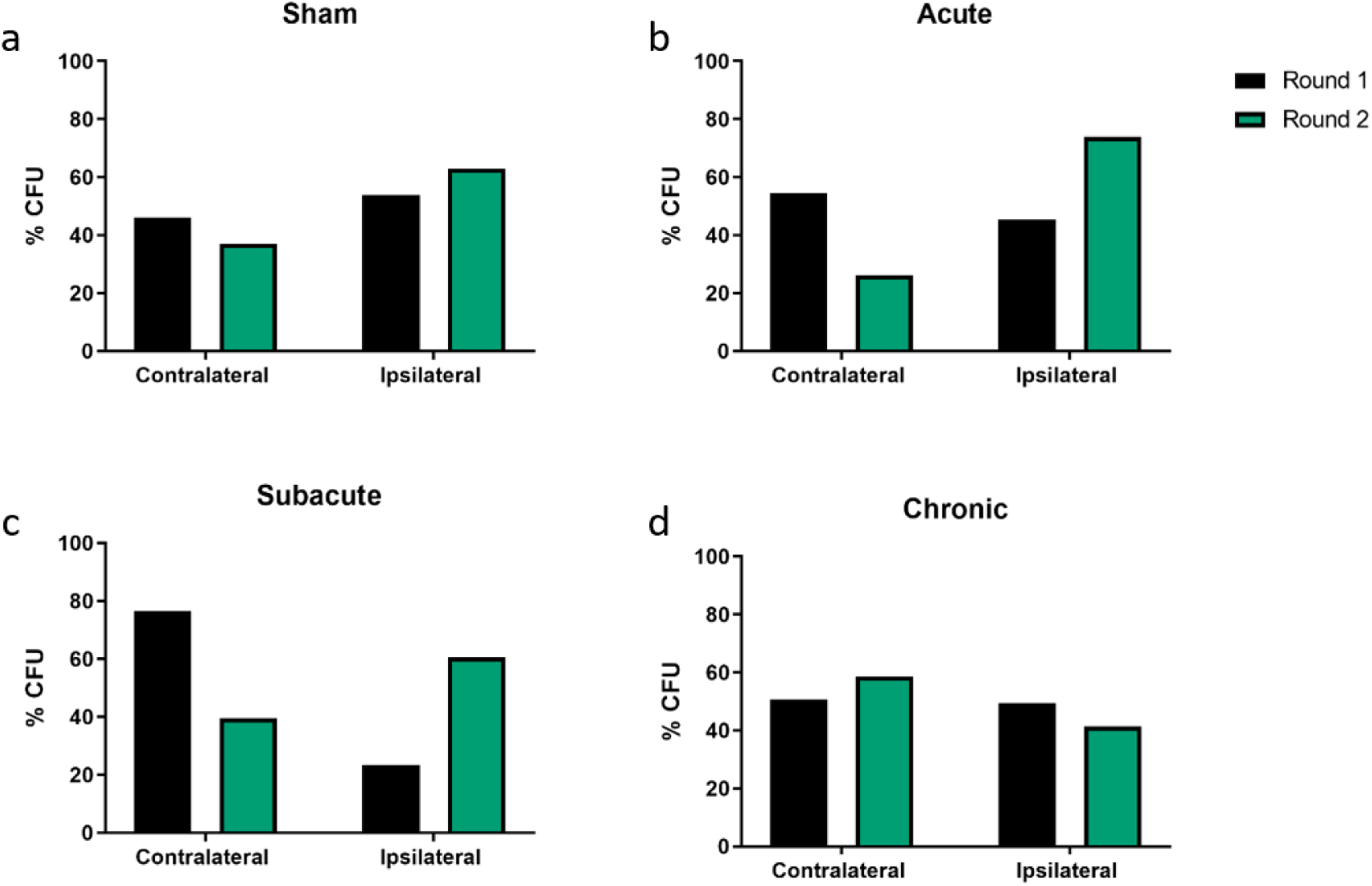
Hemisphere distribution of phage (CFU%). Recovery is quantified by % CFU for a) sham, b) acute (1dpi), c) subacute (7dpi), and d) chronic timepoints (21 dpi). Phage accumulation to the ipsilateral hemisphere increased after biopanning for both the acute and subacute timepoints. Phage distribution between hemispheres for the sham and chronic cohorts remained similar across biopanning rounds.

### NGS analysis reveals HCDR3 sequences specific to distinct injury timepoints

dAb phage libraries were sequenced via NGS; the HCDR3 sequence for each dAb was used for all subsequent analyses. This region is the only HCDR within the dAb structure that differs in canonical composition and residue length, indicating that these characteristics promote unique antigen binding specificity [21, 40]. Injury libraries yielded thousands of HCDR3s for each biopanning round, with between 200,000 to 600,000 sequences in the final biopanning round (Figure 3a). This analysis yielded a small fraction of sequences similar between timepoints, suggesting that dAb phage interacted uniquely with the neural microenvironment dependent on the temporal condition. After the final biopanning round, less than 20% of sequences from each injury library were identical with the propagation library, suggesting that injury libraries were specific to neural injury pathology (Figure 3b).

**Figure 3:**
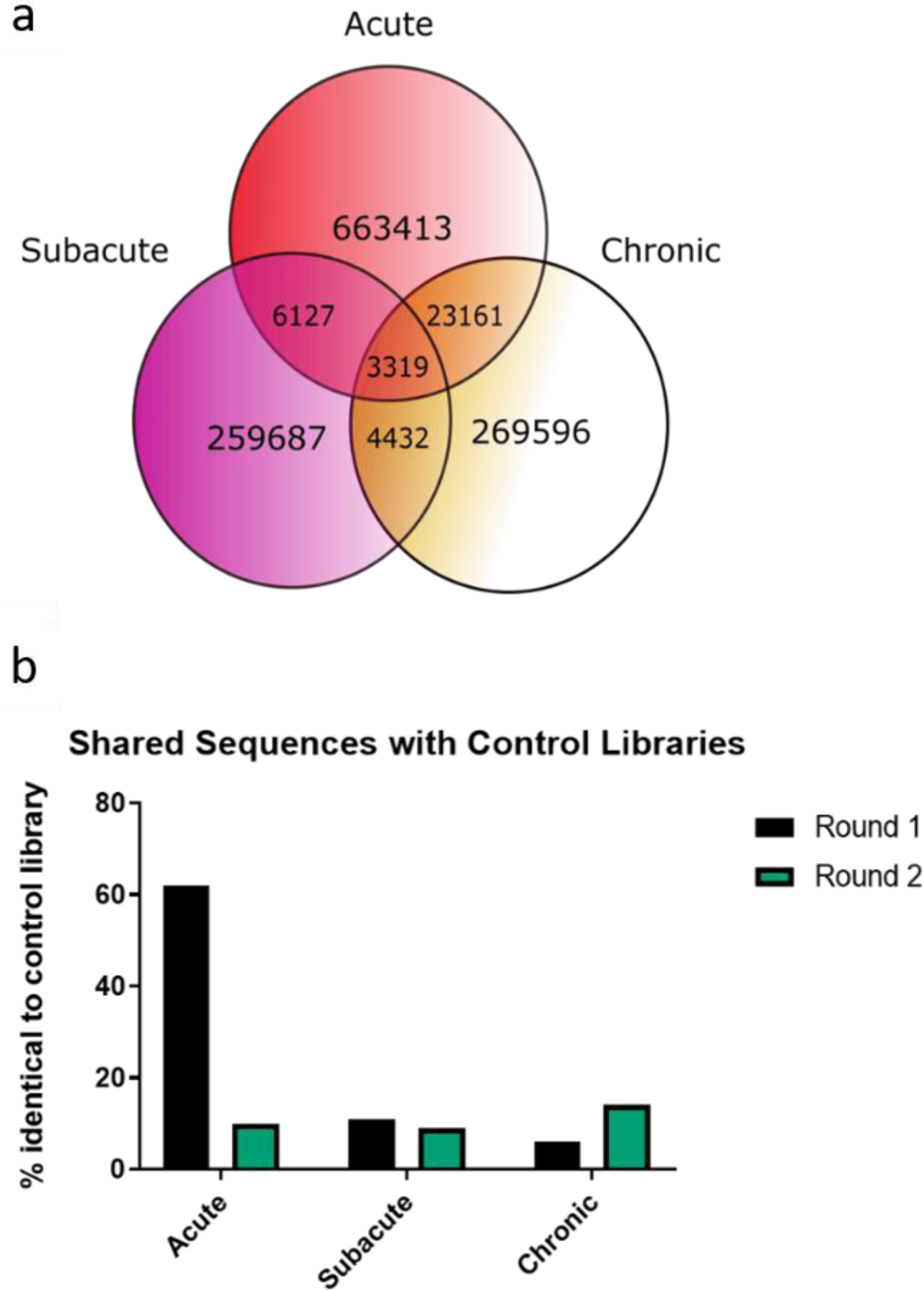
Sequence population diversity. (a) Comparison of recovered HCDR3s across injury timepoints represented by a Venn diagram. A majority of the recovered sequences were unique to their distinct timepoint, while a small fraction was found in multiple injury libraries simultaneously. (b) Comparison of recovered injury library HCDR3s against control propagation library. For the acute injury library, the percentage of sequences found in the control propagation library drastically decreased after biopanning. Both the subacute and chronic injury libraries yielded less than 20% similarity with controls across biopanning rounds.

### Biopanning increases frequency of neural injury-specific HCDR3s

Across conditions, libraries recovered from the ipsilateral hemisphere yielded substantially more sequences with higher expression (>200 reads) in the 2^nd^ biopanning round than the first (Supplementary Figure 2). This shift in frequency is representative of the biopanning process enriching the population of sequences that have preferential binding to injured neural tissue. Sequences that had an increased frequency in the final biopanning round than the first were categorized as “enriched” (Figure 4). Only 6.7% and 3.0% of sequences met this criterion for the acute and subacute libraries respectively, which provided an opportunity to target HCDR3s that were highly expressed due to affinity selection (Table 1).

**Table 1:** Percentage of HCDR3s meeting selection criteria. Enrichment is defined as an HCDR3 with increased frequency after biopanning. The z-score thresholds were 0.955 and 0.566 for acute and subacute libraries respectively, determined by the average for each injury group.

|  | Recovered | Enriched | Z-score |
| --- | --- | --- | --- |
| Acute | 663413 | 44680 (6.73%) | 500 (0.08%) |
| Subacute | 259687 | 7945 (3.06%) | 3996 (1.54%) |

**Figure 4:**
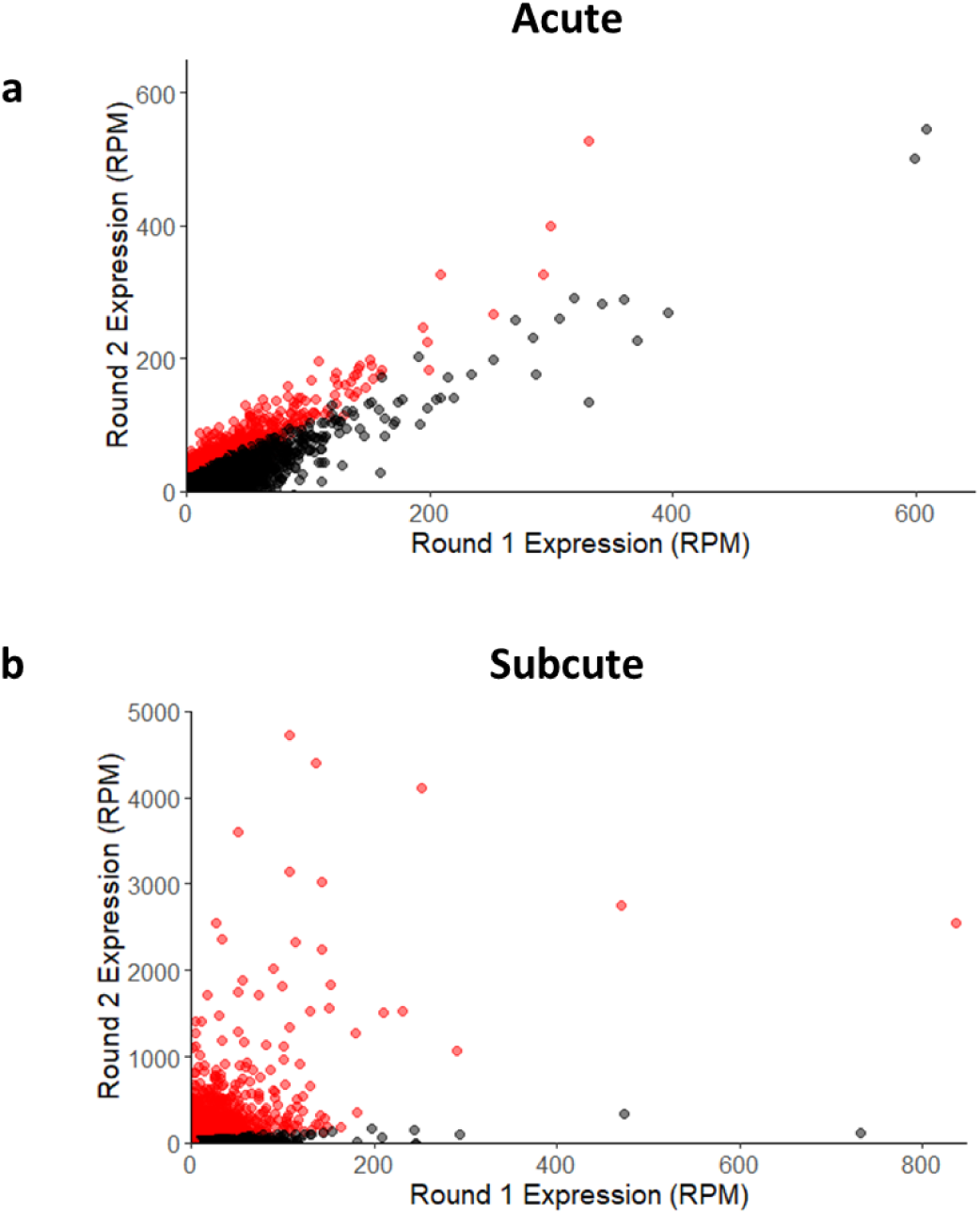
Reads per million (RPM) of sequences increased after biopanning. Relationship between individual sequence RPMs after the biopanning rounds are visualized with scatterplots for a) acute and b) subacute injury libraries. Red data points represent sequences that were enriched through biopanning.

### Phage display derived HCDR3s are temporally specific to distinct injury timepoints

Heatmaps of normalized sequence RPMS were constructed to visualize temporal relationships of the enriched HCDR3s for each timepoint. A multitude of sequences with the highest RPMS in their respective groups were observed in other timepoints post-injury (Figure 5a). For example, several HCDR3s in the acute timepoint were most highly represented in the subacute and chronic timepoints. Creating a z-score matrix of the sequences provided an opportunity to develop stringent criteria for selecting timepoint-specific sequences for HCDR3-construct design. Z-scores were averaged for each timepoint and used as a threshold to identify HCDR3s with strong preference for their distinct timepoint. Of the enriched sequences, less than 2% met z-score criteria (Figure 5b,c; Table 1). This bioinformatic analysis narrowed the pool of candidate biological motifs to an exclusive and focused group. For final selection, HCDR3s were required to 1) be unique to a distinct temporal phase post-injury, as determined by z-score normalization and comparison against other injury libraries, 2) be enriched after biopanning, and 3) not be present in control libraries. Based on these guidelines, two HCDR3s were selected for each injury group; one sequence with the highest frequency and another with the highest fold enrichment value (Table 2). Biotinylated HCDR3 peptides were synthesized and purified, validated by MALDI mass spectrometry (Figure 6).

**Table 2:**
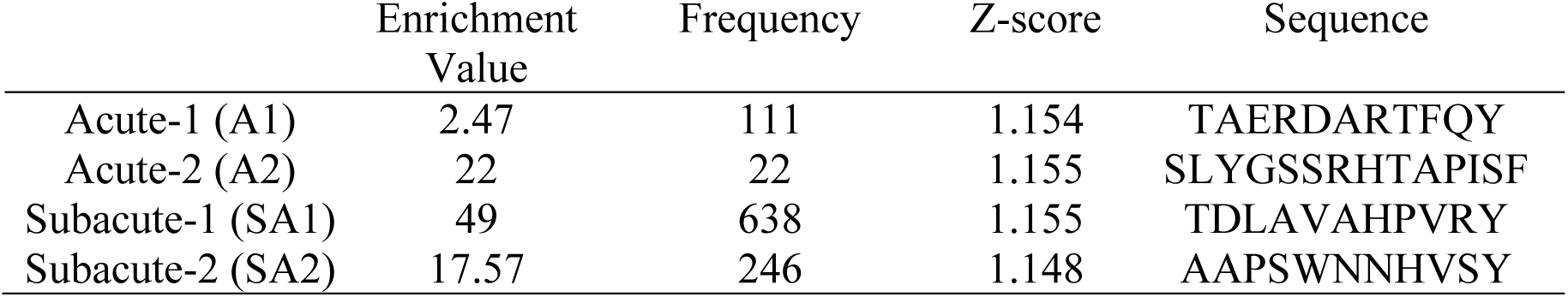
Selected HCDR3s.

**Figure 5:**
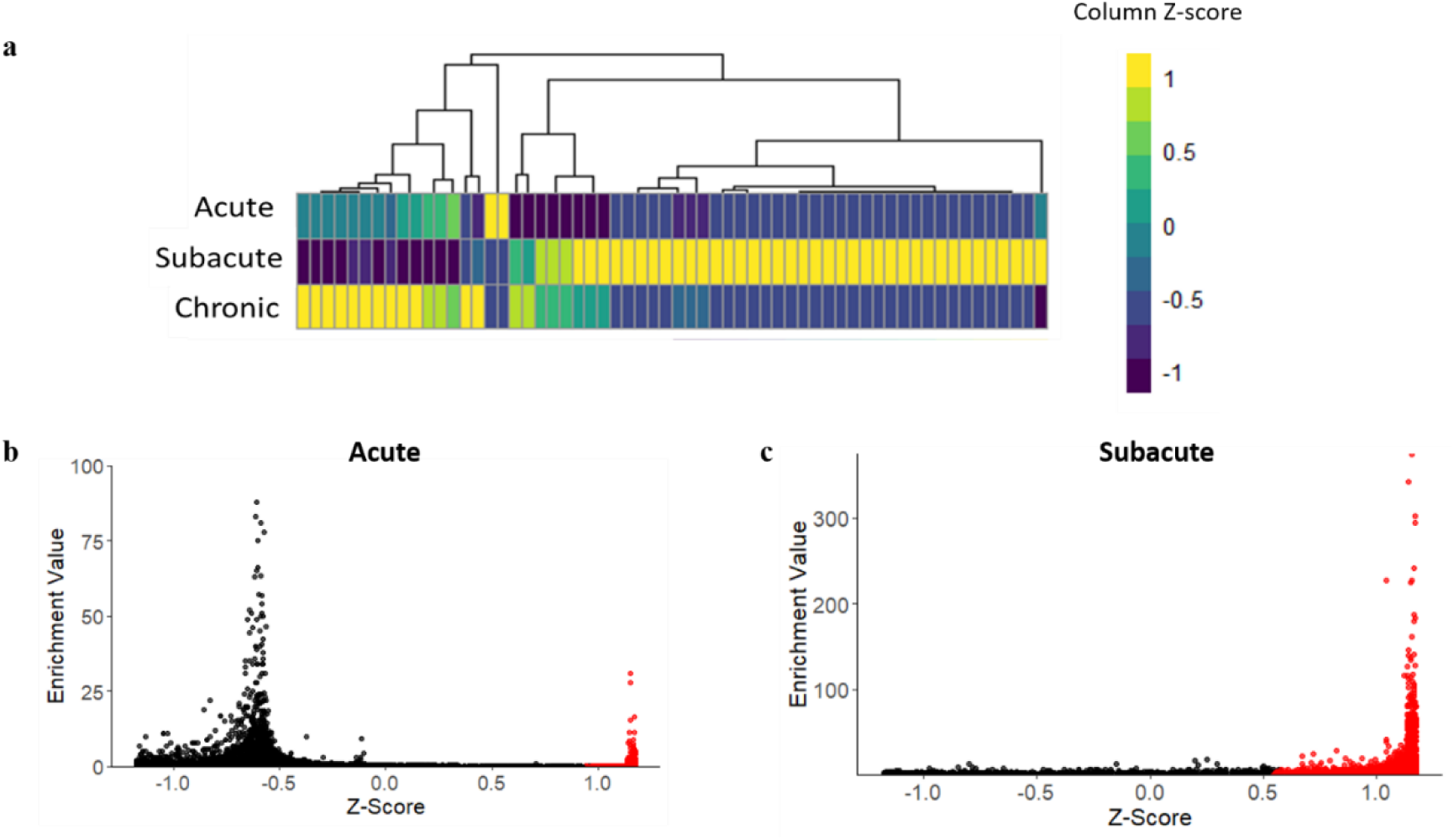
Analysis of HCDR3 temporal specificity. a) Representative heatmap of the top 20 highest frequency HCDR3s identified in each injury timepoint and their expression in adjacent timepoints. Z-scores are calculated by column (individual sequences). Scatter plots were generated to visualize the relationship between enrichment value (defined as Round 2 reads/Round 1 reads) and z-score for b) acute and c) subacute injuryHCDR3s. Red data points represent sequences that met z-score threshold criteria.

**Figure 6:**
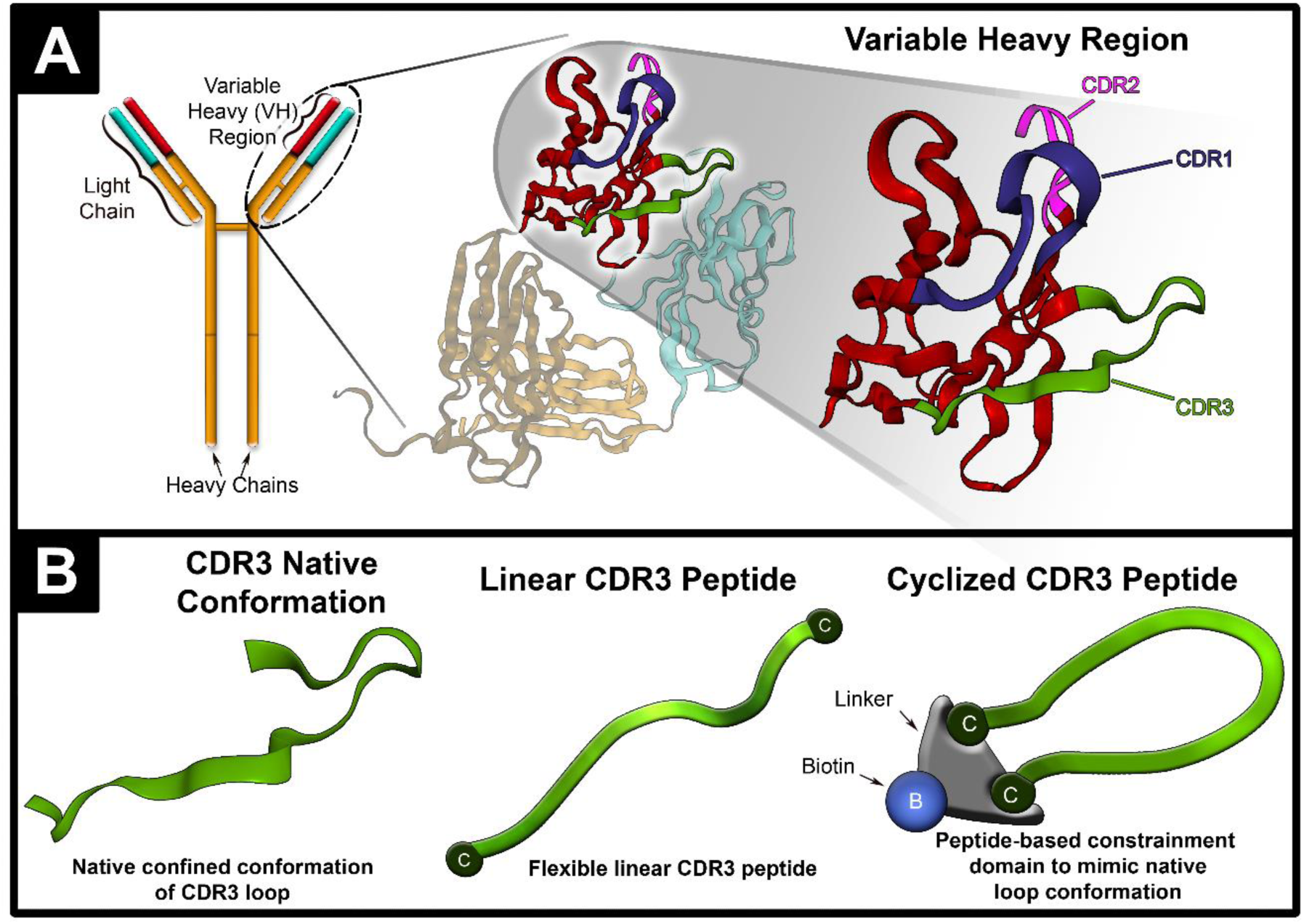
Cyclized HCDR3 peptides. a) Representation of heavy complementarity determining regions (HCDRs). Domain antibody (dAb) consist of the variable heavy region of a full length antibody. Each dAb contains three HCDRs, with the HCDR3 facilitating antigen binding. b) Representation of cyclized HCDR3 peptide. Each HCDR3 peptide construct is conjugated to a bivalent peptide linker that contained a C-terminal biotin.

### Validation of spatiotemporal affinity

Immunohistochemistry (IHC) was used to evaluate the recognition of HCDR3-constructs to CCI injury sections. IHC with acute-1 construct (A1) yielded no detectable signal on injured tissue. Acute-2 construct (A2) showed significant bioreactivity determined by the fluorescence on 1 dpi tissue compared to sham tissue (p= 0.0120) and 7 dpi tissue (p = 0.0221). While trending toward significance, no statistically significant differences were observed between 1 dpi and 21 dpi tissue (p = 0.0658). No detectable signal was observed with subacute-2 construct (SA2), but positive stain with (SA1) was observed in the peri-injury region of the 7 days post-injury tissue, while this localization was not observed in sham brain sections (p = 0.0079) (Figure 7). No significant differences were observed between SA1 bioreactivity on 7 dpi and 1 dpi tissue (p = 0.0993) or 21 dpi tissue (p = 0.0780). Control constructs (derived from spleen, heart, and propagation phage library; Supplementary Table 3) showed no detectable signal on injured tissue at 1 or 7 dpi, demonstrating that the positive signal we observed from the A2 and SA1 were not due to non-specific artifact derived from construct structure (Supplementary Figure 3).

**Figure 7:**
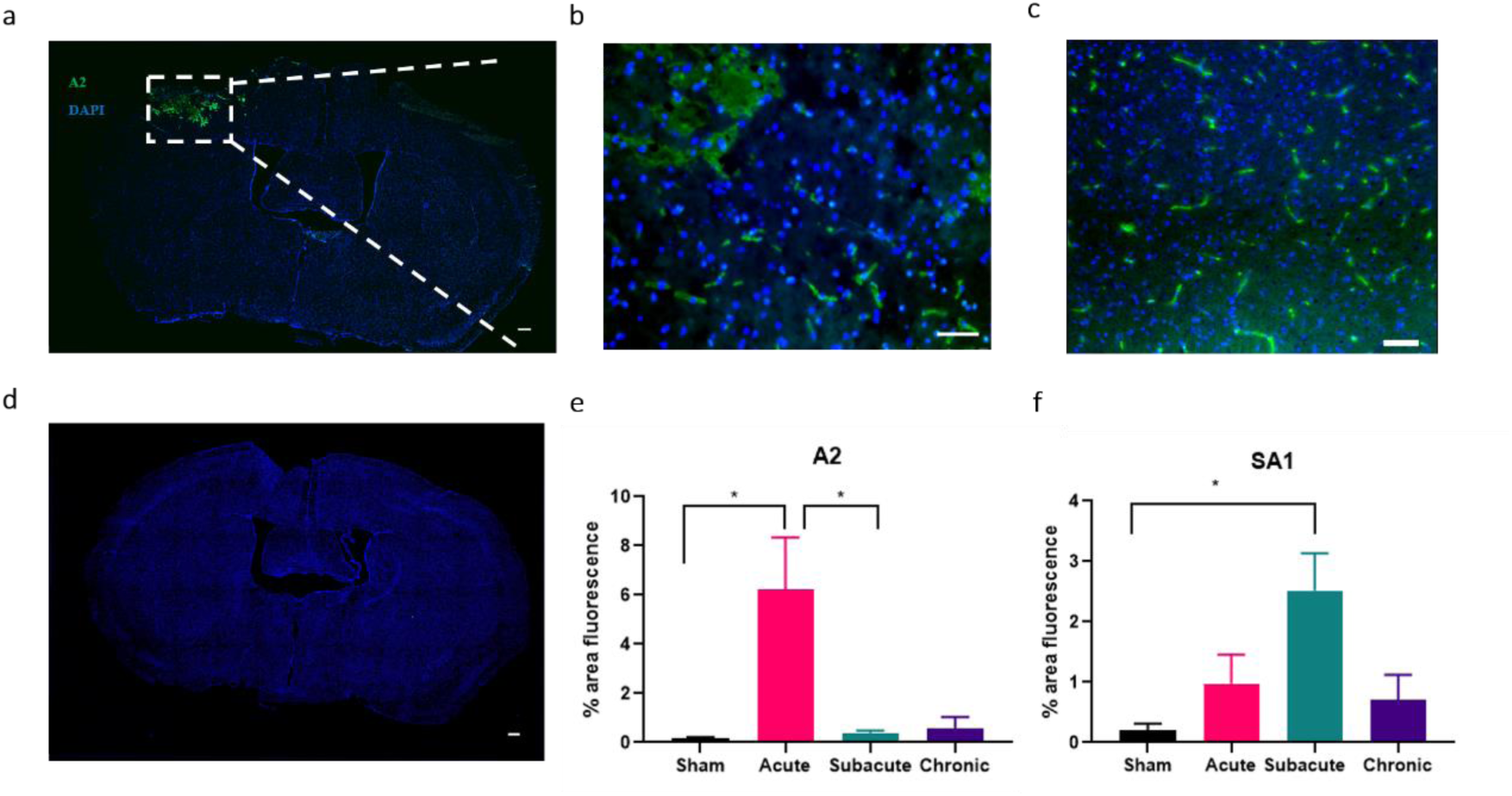
HCDR3 constructs show selectivity to injured tissue. a) Qualitative representation of injury-specific HCDR3 (green) and cell nuclei (blue) in 1 dpi tissue. ROI represented in white box. Scale bar = 200 µm. b) 20x magnification of A2 construct staining on 1 dpi tissue. Scale bar = 100 µm. c) 20x magnification of SA1 construct staining on 7 dpi tissue. Scale bar = 100 µm. d) qualitative representation of sham control. Scale bar = 200 µm. e and f) Quantification of % area fluorescence in 1500 µm x 1500 µm ROI (n=5-6). Data expressed in mean + SEM. * p < 0.05.

### IP-MS isolates proteins involved in mitochondrial dysfunction and neurodegenerative processes

IP-MS analysis identified 18 and 20 proteins specific to the injury condition when using A2 and SA1 as capture antibodies, respectively (FDR <0.01) (Supplementary Table 4, 5). Ontological analysis of candidate proteins revealed several biological processes that were dominant and similarly represented across groups, such as metabolic and cellular processes (Figure 8). Only SA1 isolated proteins linked to behavioral and developmental processes (5.6%) that were not represented in the A2 condition (Figure 8). Pathway analysis of A2-specific proteins identified the TCA cycle and pyruvate metabolism pathway as unique and highly represented processes (20% and 13% respectively) (Figure 9). Comparatively, proteins implicated in Parkinson’s disease and apoptosis signaling pathways were highly represented for SA1-specific proteins (both 11%) (Figure 9).

**Figure 8:**
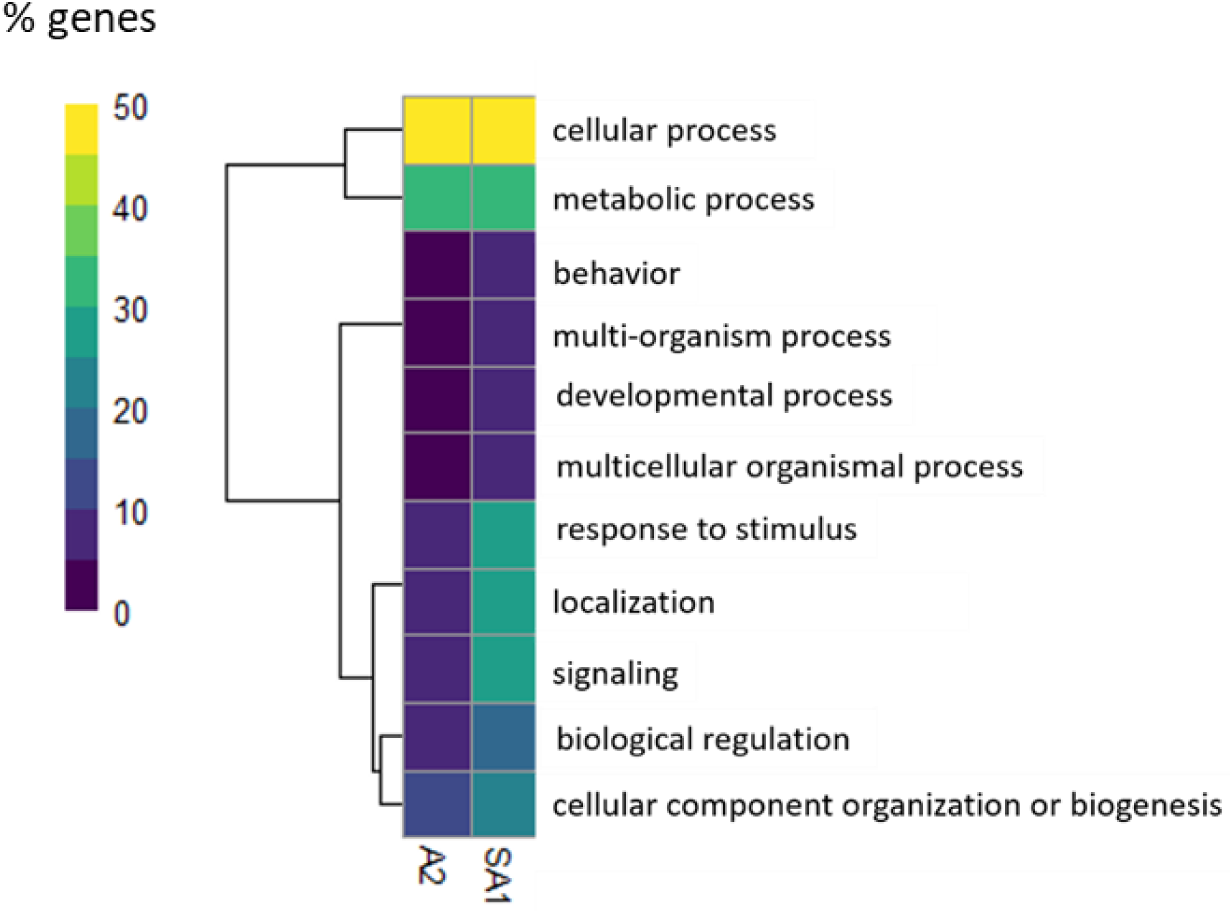
Biological process categorization of identified proteins isolated by A2 and SA1. Categories hierarchically clustered by percent of proteins identified in biological processes. A2 and SA1 had similar distribution of proteins involved in cellular and metabolic processes. Comparatively, proteins isolated by SA1 also included pathways associated with localization, response to stimulus, and signaling than A2-associated proteins.

**Figure 9:**
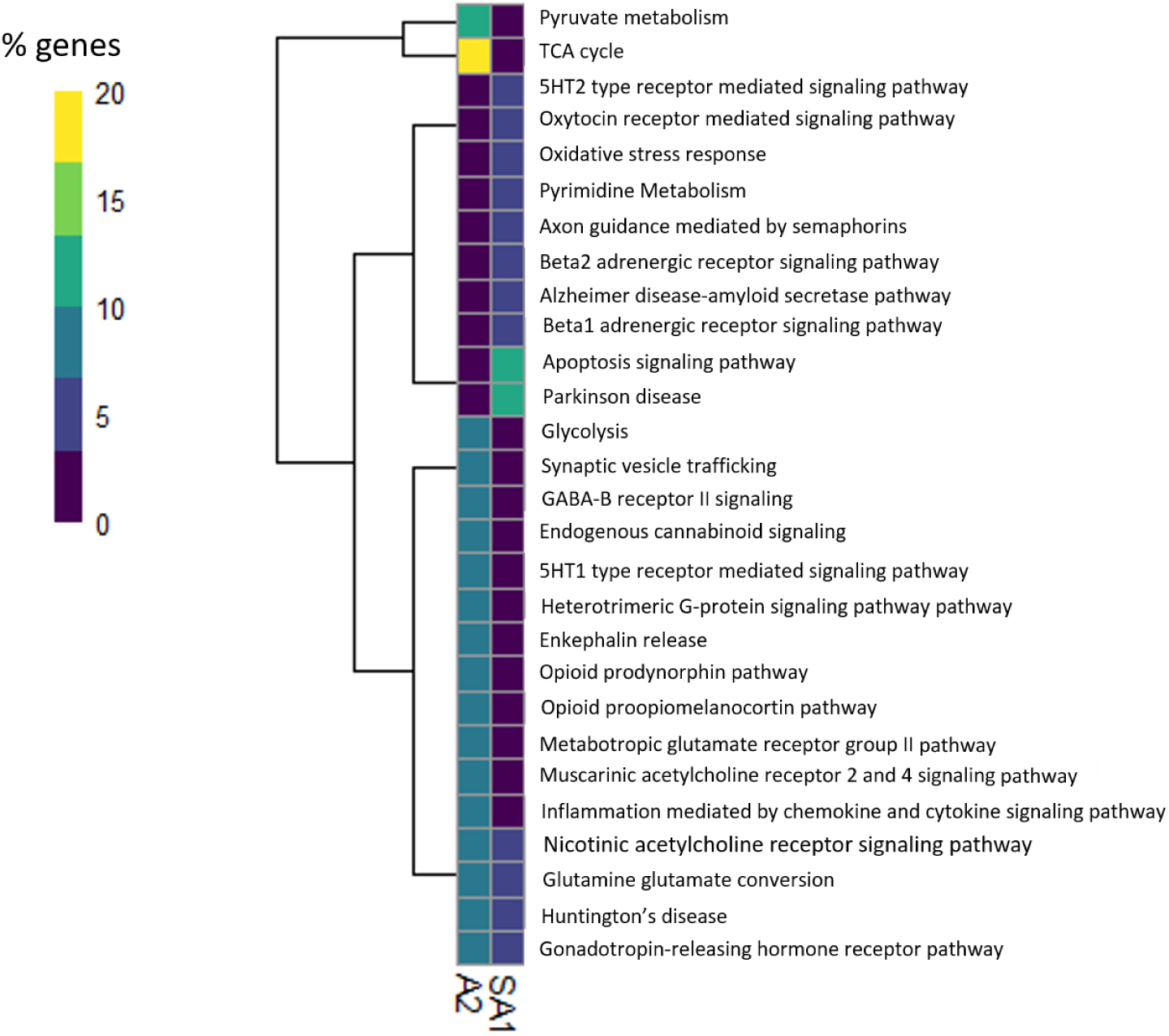
Pathway categorization of identified proteins isolated by A2 and SA1. Categories hierarchically clustered by percentage of proteins identified in pathway analysis. Proteins identified in the A2 condition were highly expressed in the TCA cycle and pyruvate metabolism in comparison to other pathway analysis categories. Proteins identified in the SA1 condition were highly expressed in Parkinson’s disease and apoptotic signaling pathways by comparison.

Identified proteins with the highest number of peptides recovered by mass spectrometry analysis were highly involved in the categorized pathways (Table 3). Citrate synthase (CS), and succinyl CoA synthetase subunit β were identified as prominent components of the TCA cycle, while CS was also represented in the pyruvate metabolism pathway. Heat shock cognate 71 kDa and endoplasmic reticulum chaperone binding immunoglobulin protein (ER chaperone BiP) were identified as components of both the Parkinson’s disease and apoptosis signaling pathways. The high volume of peptides recovered by MS and their involvement in highly represented pathways suggest that these proteins are probable targets of the HCDR3 constructs.

**Table 3:** Candidate proteins isolated by HCDR3 constructs.

| Targeting construct | Description | Accession | # Peptides |
| --- | --- | --- | --- |
| A2 | Succinate--CoA<br>ligase [ADP-forming] subunit<br>beta, mitochondrial | Q9Z2I9 | 4 |
|  | citrate synthase,<br>mitochondrial | Q9CZU6 | 3 |
| SA1 | Heat shock cognate<br>71 kDa protein | P63017 | 6 |
|  | Endoplasmic<br>reticulum chaperone<br>BiP | P20029 | 6 |

## DISCUSSSION

A confounding factor in the diagnosis and treatment of TBI is temporal complexity of pathological progression. Analysis of temporal biomolecular mechanisms provide insight on injury progression that may better inform the development of theranostic tools. In this study, we leveraged the power of unbiased phage display to identify and develop novel biomolecular motifs that specifically recognize elements of acute and subacute TBI pathology. *In vivo* phage biopanning was conducted in a CCI rodent model at three distinct time points following injury (1, 7, and 21 dpi) to perform a robust bioinformatics driven assessment of enriched phage populations for each timepoint. The spatiotemporal specificity of HDCR3-constructs based on the NGS data were validated first with IHC analysis, demonstrating the strength of this high throughput screening and sequencing methodology. Using IP-MS, we positively identified several candidate proteins involved in metabolic and neurodegenerative processes as potential targets for A2 and SA1, respectively. The elegance of phage-based approach in contrast to traditional neuroproteomics (i.e., whole brain tissue analysis to identify differentially expressed proteins via mass spectrometry [11]), is that phage biopanning leverages molecular evolution to narrow the molecular pool for biomarker candidate selection.

dAb libraries are advantageous to screening against neural tissue *in vivo* due to their small size (12-15 kDa), high affinity, and ability to effectively bind to brain vasculature under normal BBB conditions [24, 41]. Interestingly, dAb phage accumulation in naïve and chronic injury neural tissue was comparable across biopanning rounds while accumulation within acute and subacute injury groups drastically increased from round 1 to 2. While phage displayed motifs can target brain vasculature, *in vivo* screening in models of injury and degeneration have demonstrated that BBB disruption provides opportunity for extravascular binding of the phage [42, 43]. Notably, the disrupted state of the BBB after disruption persists in the CCI model up to 7 dpi, allowing macromolecules and potential therapeutic agents to gain extravascular access via intravenous circulation [44, 45]. The compromised condition of the BBB thus may have permitted intravenously injected phage to access extravascular targets; our data corroborate this phenomena whereby relatively lower percentage of phage accumulation in sham and chronic injury cohorts (Figure 2) [18, 45].

Recent sequencing advancements in NGS capabilities are instrumental to the identification of candidate biological motifs in phage display libraries. NGS analysis provides an opportunity to uncover the entire population of phage display libraries at a sequencing space of 10^5^-10^7^ in comparison to 20-100 for traditional Sanger sequencing methods [46]. This analysis also minimizes the probability of selecting false positive clones that may be overrepresented in the library due to propagation advantages, amplified as a consequence of repetitive screening [47, 48]. Applying NGS analysis to the screening results circumvented this dilemma by providing a platform to achieve a thorough analysis with only two biopanning rounds, thereby overcoming a large drawback of utilizing phage display technology. Both advantages are critical for the analysis of a library derived from *in vivo* biopanning of the neural injury microenvironment.

In lieu of recombinant protein production, we addressed the challenge of bioreactivity validation by designing novel peptide-based HCDR3 constructs that mimic the constrained HCDR loop structure, motivated by prior studies [20], thereby enabling high-throughput production via direct peptide synthesis and facile biochemical modifications to fabricate the constrained cyclic HCDR3 loop structure. The HCDR3 has been identified as the main contributor to binding specificity of antibodies and truncated antibody fragments. Prior studies have highlighted the utility of generating HCDR3 peptide variants as a “synthetic antibody” with comparable binding efficiency to full length antibodies [20, 31]. Our data further supports this reductionist approach and future studies will focus on potential mechanisms and molecular tuning to optimize the HCDR loop structure.

Our validation results readily demonstrated the critical need for thorough testing of each phage identified candidate motif. Most prominent, A1 was identified based on our selection criteria for the acute timepoint, namely high frequency in biopanning round 2, yet IHC assessment did not show detectable bioreactivity with fixed mouse brain tissue at 1 dpi. In contrast, A2, selected namely for the high fold enrichment value from biopanning round 1 to round 2, showed high sensitivity and affinity to the peri-injury region at 1dpi (Figure 6). For the subacute constructs, the opposite effect was observed with constructs targeting subacute injury, with SA1, a construct selected for its high frequency, positively binding to injured neural tissue. A1 had the highest observed frequency for its timepoint, yet it only exhibited a fold-enrichment value of 2.47; much lower than A2’s enrichment value of 22. Furthermore, SA2’s enrichment value of 17.57 dwarfed in comparison to SA1’s value of 49 (Table 2). These results may suggest that enrichment facilitated by biopanning plays a critical role in the ability of the HCDR3 to bind successfully to its target. However, more testing is required to fully understand the factors that influence bioreactivity of a phage-selected motif.

A2 positive staining on 1 dpi tissue showed significant bioreactivity compared to sham and subacute tissue, with a concentrated visual appearance restricted to the injury penumbra. In comparison, SA1 positive staining on 7 dpi tissue was punctate and more dispersed, consequently yielding a lower average fluorescence per area than the acute group. While A2 demonstrated nearly exclusive bioreactivity to acute injury, SA1 exhibited a gradual trend in bioreactivity over time although bioinformatic analysis suggested this construct was specific to subacute injury alone. Positive staining on 7dpi was statistically significant in comparison to the sham cohort, but a notable increase is observed on 1 dpi tissue before peaking at 7dpi, then tapering at 21 dpi. This result may indicate targeting of antigens that are expressed at a minimal level acutely after injury but increase in expression over time.

Further validation of our HCDR3 constructs was performed by IP-MS analysis. We identified potential acute TBI pathology targets of A2 as critical metabolic processes mediators. Pyruvate metabolism and TCA cycle, two pathways revealed by subsequent A2 target pathway analysis, work in tandem to regulate cerebral metabolism [49]. After TBI, these pathways are inhibited due to oxidative stress damage caused by mitochondrial dysfunction [50]. Ontological analysis revealed two individual components targeted by A2 that are implicated in these pathways and highly represented in the mass spectrometry data; succinate CoA ligase β and citrate synthase (CS) (Table 3). Alterations in succinate-CoA ligase β cause mitochondrial dysfunction and negatively impacts the central nervous system with disorders such as encephalomyopathy [51, 52]. This subunit is increased in the rat brain proteome three hours after hemorrhagic stroke in comparison to naïve controls, providing evidence for time-dependent upregulation after neural injury [14]. Interestingly, CS is significantly downregulated in comparison to controls acutely after diffuse axonal injury and CCI [53, 54]. However, CS expression may be dependent on both severity and time, with significantly decreased expression of CS in severe weight drop models at 6, 24, 48, and 120 hours post-injury in comparison to mild TBI conditions [55]. The isolation of this enzyme by A2 is possibly due to its interaction with succinate-CoA and pyruvate dehydrogenase. Each protein is essential to the functionality of the TCA cycle, and both succinate-CoA and pyruvate dehydrogenase actively inhibit CS [56]. While A2 isolated several proteins involved in the TCA cycle, it is probable that CS was immunoprecipitated along with other TCA cycle proteins that are highly upregulated after injury.

SA1 isolated proteins strongly associated with neurodegenerative processes such as Huntington’s, Alzheimer’s, and Parkinson’s disease. (Figure 8). Heat shock cognate 71 kDa and ER chaperone BiP, members of the heat shock protein 70 family, were both identified as components of Parkinson’s disease and the apoptosis signaling pathway. ER chaperone BiP, a monitor of endoplasmic reticulum stress is induced in Alzheimer’s disease in response to protein misfolding and cell death [57, 58]. Additionally, a reduction in ER chaperone BiP expression leads to the acceleration of prion disease pathology [59]. Comparatively, an increase in ER chaperone BIP expression is suggested to be neuroprotective in models of brain ischemia [60, 61]. Recent studies suggest that heat shock cognate 71 kDa, a cytosolic facilitator of protein folding and degradation, may have a strong interaction with Tau protein, a hallmark of Alzheimer’s disease [62]. This protein has been suggested as a possible therapeutic target for stroke and TBI as well, as its overexpression may reduce apoptosis and inflammation [63]. TBI is a risk factor for neurodegenerative diseases, and many factors in the secondary injury cascade run parallel to degenerative pathology such as neuronal cell death [64, 65].

The isolation of proteins implicated in mitochondrial dysfunction and neurodegeneration 1 and 7 dpi respectively reflects the influence injury pathology has on neurodegenerative diseases. For example, mitochondrial dysfunction is a critical early-occurring factor that influences apoptosis and reactive oxygen species overproduction in both Alzheimer’s patients and animal models [66, 67]. This contribution to toxic neurodegenerative pathology is not restricted to Alzheimer’s disease, but is also observed in Parkinson’s disease, Huntington’s disease, and amyotrophic lateral sclerosis [52, 68, 69]. Despite injury and neurodegenerative pathology sharing mitochondrial dysfunction as an attribute, the connection between TBI and toxic mechanisms of neurodegenerative diseases, along with its potential therapeutic value, has yet to be fully elucidated [65, 70]. The unbiased identification of proteins involved in these processes acutely and subacutely in TBI through phage display therefore provides insightful perspective on the link between brain injury and acquisition of neurodegenerative diseases.

## LIMITATIONS

While the current study successfully explores the utility of phage display-facilitated elucidation of TBI, there are limitations to the presented findings. First, the study implements a one-time CCI as the chosen TBI animal model, which cannot recapitulate the effects of more complex instances of TBI, such as repetitive head injury. However, no one animal model can fully replicate the human TBI experience [8]. Therefore, it would be ideal to apply this phage display discovery pipeline to various animal models such as fluid percussion injury and blast injury to compare and validate results.

Additionally, cognitive and behavioral measures were not considered during the study, such as stress response. Stress affects the outcome of TBI by inducing inflammation and consequently influencing neuropathological changes that cause prolonged anxiety- and depression-like behaviors [71]. TBI has been linked with post-traumatic stress disorder, which is attributed with changes in neurocircuitry such as the hippocampus-striatum pathway [72–74]. Assessing stress response in a future study would bring a new dimension to this work by providing information on behavioral stressors effects on acute injury pathology, and how these changes influence the neural milieu over time. This multi-dimensional approach would greatly contribute to the elucidation of TBI behavioral effects while also insight on potential therapeutic targets.

## CONCLUSION

The current study not only identified proteins specific to temporal brain injury phases, but simultaneously developed targeting constructs for these candidates. The design of HCDR3 constructs that specifically bind to acute and subacute injury provides a foundation for the development of theranostic tools. This discovery will allow for future characterization of the candidate targets through several conditions within the neural injury microenvironment, in addition to the refinement of HCDR3-constructs as a targeting modality to detect and treat TBI.

## ACKNOWLEDGEMENTS

We acknowledge the following facilities and/or personnel for support: ASU Genomics Facility and Dr. Shanshan Yang for next generation sequencing support, ASU Mass Spectrometry Core and Dr. Timothy Karr for mass spectrometry services, Amanda Witten for figure illustrations, and Nathan Borgogni for immunohistochemistry assistance. We also acknowledge the following funding sources: NIH NICHD 1DP2HD084067 and 1DP2HD084067-01S1 (SES), NIH NINDS 1R21NS107985 (SES), ARCS Scholar (BIM), and ASU Biomimicry Seed funding (CD, NS, SES).

