## Supplemental Data for "Uncovering temporally sensitive targeting motifs for traumatic brain injury via phage display"

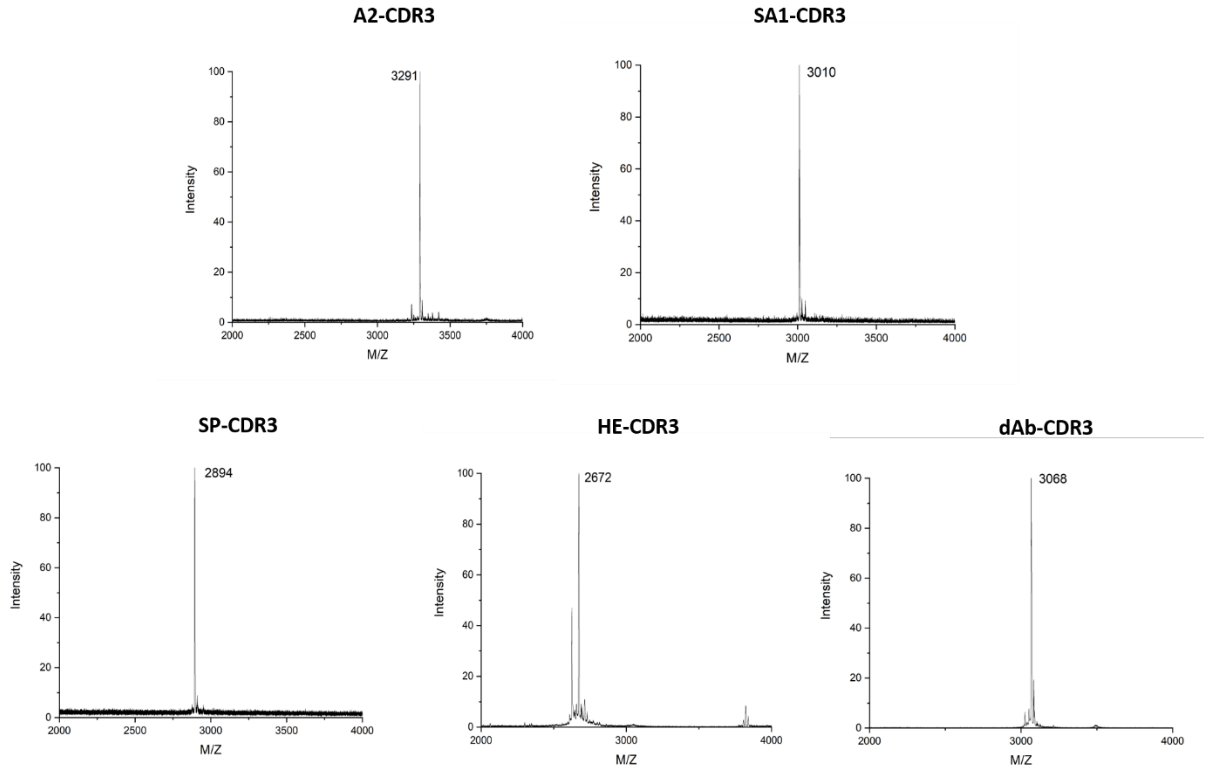


**Supplementary Figure 1: MALDI MS of synthesized HCDR3 cyclic peptides after HPLC purification**

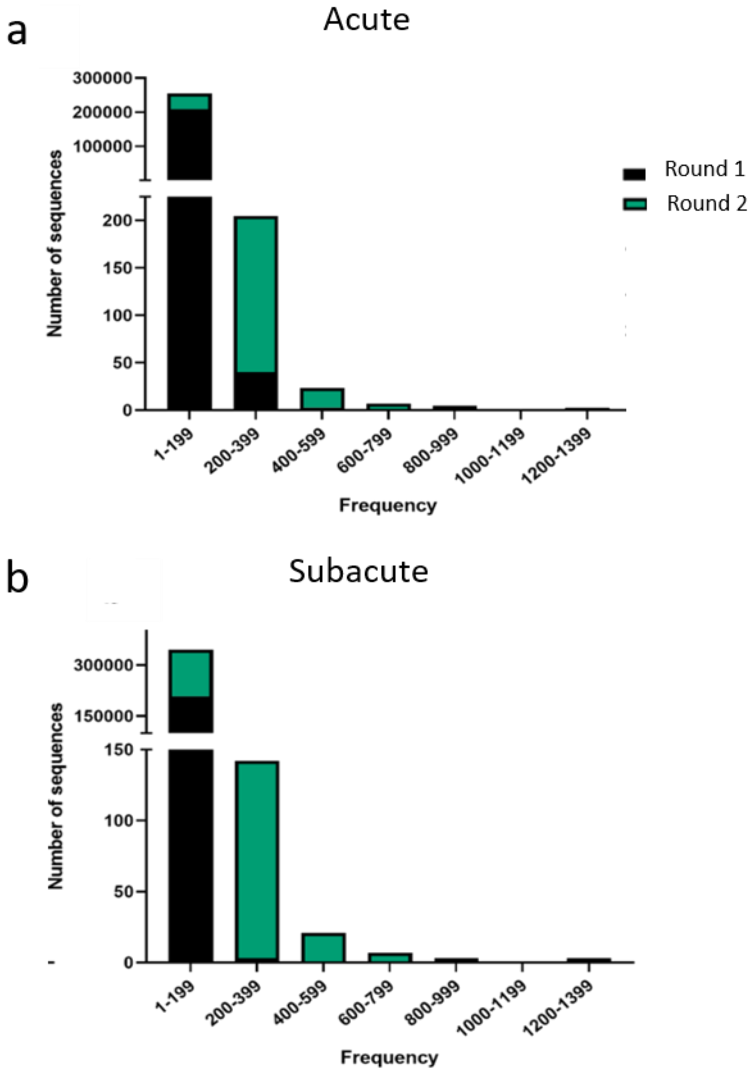


**Supplementary Figure 2: Frequency distribution of recovered HCDR3s within injury libraries.** Round 2 yielded more sequences in higher ranges (>200 reads) than after round 1 of biopanning. This shift in frequency is representative of the biopanning process enriching the population of ipsilateral-specific sequences.

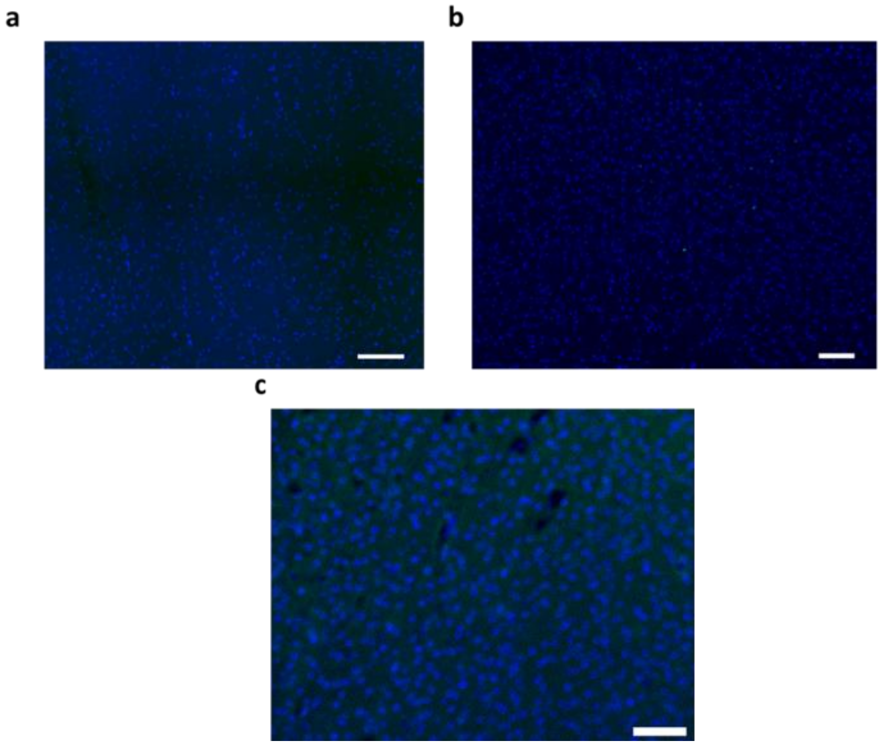


**Supplementary Figure 3: Control constructs show no detectable bioreactivity on injured neural tissue.** Constructs were designed based on sequences highly expressed in control libraries. a) Spleen, b) dAb propagation and c) heart HCDR3 constructs showed no detectable bioreactivity with injured neural tissue. Scale bar = 100 µm.

**Supplementary Table 1: dAb sequencing primers for MiSeq 2 x 300 bp module.** Underlined portion indicates Illumina overhang adapter sequence.

| Primer Name | Sequence |
| --- | --- |
| dAb For | 5' TCGTCGGCAGCGTCAGATGTGTATAAGAGACAGCAGCTGTTGGAGTCTGGGG 3' |
| dAb rev | 5' GTCTCGTGGGCTCGGAGATGTGTATAAGAGACAGGAGACGGTGACCAGGGTTC 3' |

**Supplementary Table 2: Phage accumulation in ipsilateral and contralateral hemispheres determined by CFU/g**

| | | Sham | Acute | Subacute | Chronic |
|---|---|---|---|---|---|
| Round 1 (CFU/g) | Heart | $5.54 \times 10^5$ | $4.54 \times 10^4$ | $5.54 \times 10^5$ | $1.68 \times 10^6$ |
| | Spleen | $9.50 \times 10^6$ | $1.05 \times 10^7$ | $3.83 \times 10^6$ | $3.57 \times 10^6$ |
| | Contralateral | $1.08 \times 10^5$ | $3.92 \times 10^3$ | $1.21 \times 10^6$ | $5.69 \times 10^4$ |
| | Ipsilateral | $9.87 \times 10^4$ | $1.73 \times 10^3$ | $5.13 \times 10^5$ | $7.94 \times 10^4$ |
| Round 2 (CFU/g) | Heart | $1.90 \times 10^6$ | $4.79 \times 10^6$ | $1.17 \times 10^4$ | $8.72 \times 10^6$ |
| | Spleen | $3.11 \times 10^6$ | $1.77 \times 10^7$ | $1.867 \times 10^7$ | $9.20 \times 10^7$ |
| | Contralateral | $7.07 \times 10^3$ | $3.92 \times 10^3$ | $1.15 \times 10^4$ | $3.58 \times 10^4$ |
| | Ipsilateral | $1.54 \times 10^4$ | $7.69 \times 10^5$ | $2.26 \times 10^4$ | $2.52 \times 10^4$ |

**Supplementary Table 3: HCDR3 control constructs.** HE-CDR3 = heart, SP-CDR3 = spleen, dAb-CDR3 = dAb propagation.

| HCDR3-Construct | Sequence |
|---|---|
| HE-CDR3 | TGHEGENEMAS |
| SP-CDR3 | GPLDGKEEELRF |
| dAb-CDR3 | GGDTFRDASQSMHF |

**Supplementary Table 4: A2 isolated proteins determined by mass spectrometry (FDR<0.01)**

| Accession | Description |
|---|---|
| P18872 | Guanine nucleotide-binding protein G(O) subunit alpha |
| P51863 | V-type proton ATPase subunit d |
| Q62277 | Synaptophysin |
| Q8BG05 | Heterogeneous nuclear ribonucleoprotein A3 |
| Q8R010 | aminoacyl tRNA synthase complex-interacting multifunctional protein 2 |
| P15105 | Glutamine synthetase |
| P35486 | Pyruvate dehydrogenase E1 component subunit alpha, somatic form, mitochondrial |
| P42669 | Transcriptional activator protein Pur-alpha |
| P61164 | Alpha-centractin |
| Q6ZQ38 | cullin-associated nedd8-dissociated protein 1 |
| Q8BG05 | Heterogeneous nuclear ribonucleoprotein A3 |
| Q8VHF2-1 | Cadherin-related family member 5 |
| Q9WV02-1 | RNA-binding motif protein, X chromosome |
| Q9Z2I9 | Succinate--CoA ligase [ADP-forming] subunit beta, mitochondrial |
| Q9WUM5 | Succinate--CoA ligase [ADP/GDP-forming] subunit alpha, mitochondrial |
| Q9CZU6 | citrate synthase, mitochondrial |
| P16330 | 2',3'-cyclic-nucleotide 3'-phosphodiesterase |

**Supplementary Table 5: SA1 isolated proteins determined by mass spectrometry (FDR<0.01)**

| Accession | Description |
|---|---|
| Q8C0N1 | Kinesin-like protein KIF2B |
| Q3U6U5 | Putative GTP-binding protein 6 |
| P62918 | 60S ribosomal protein L8 |
| P63017 | Heat shock cognate 71 kDa protein |
| P20029 | 78 kDa glucose-regulated protein |
| P19246 | Neurofilament heavy polypeptide |
| P15105 | Glutamine synthetase |
| Q6P5F9 | Exportin-1 |
| P62754 | 40S Ribosomal Protein S6 |
| P04370-4 | Isoform 4 of Myelin basic protein |
| O08553 | Dihydropyrimidinase-related protein 2 |
| Q9CXW4 | 60S ribosomal protein L11 |
| P68254-1 | 14-3-3 protein theta |
| Q8BZ36 | RAD50-interacting protein 1 |
| Q7TQH7 | Low-density lipoprotein receptor-related protein 10 |
| Q99246 | Voltage-dependent L-type calcium channel subunit alpha-1D |
| Q9DBB1 | Dual specificity protein phosphatase 6 |
| Q8CHC4 | Synaptojanin-1 |
